# Reward prediction error neurons implement an efficient code for reward

**DOI:** 10.1101/2022.11.03.515104

**Authors:** Heiko H. Schütt, Dongjae Kim, Wei Ji Ma

## Abstract

We use efficient coding principles borrowed from sensory neuroscience to derive the optimal population of neurons to encode rewards from a probability distribution. We find that the response properties of dopaminergic reward prediction error neurons in a rodent and a primate data set are similar to those of the efficient code in many ways: the neurons have a broad distribution of midpoints covering the reward distribution; neurons with higher thresholds have higher gains, more convex tuning functions, and lower slopes; moreover, their slope is higher when the reward distribution is narrower. Furthermore, we derive learning rules that converge to this efficient code; the learning rule for the position of the neuron on the reward axis closely resembles the learning rule of distributional reinforcement learning. Thus, reward prediction error neuron responses may be optimized to broadcast an efficient reward signal, forming a connection between efficient coding and reinforcement learning, two of the most successful theories in computational neuroscience.

## Main

Processing rewards is critical for much of cognition, including decision making, planning, and learning. An important reward representation in the brain is maintained by reward prediction error neurons (RPENs) [1]. These dopaminergic neurons in the midbrain respond to received rewards relative to an expectation based on past experience. The existence of these neurons is the most prominent evidence in favor of reinforcement learning (RL) in the brain [2]. Moreover, RPENs have been implied in a broad range of tasks that require value-based cognition [3]. Thus, the encoding of reward by RPENs is a cornerstone of our understanding of reward signals in the brain.

In a different domain of neuroscience, sensory processing, it has been hypothesized that neuronal codes are optimized for efficiency, i.e. to convey as much information as possible with given budgets for the number of neurons and the number of action potentials [4, 5]. The efficient coding hypothesis has long been used to account for the response properties of sensory neurons (e.g. [6–8]). Gradually, the notion of efficient coding has also made inroads in the domain of reward. Efficient coding has been invoked to explain reward-based choices [9–11], and contextual modulation of the responses of cortical reward neurons has been interpreted as a form of efficient coding [12]. In a time interval estimation task, an RL agent that encoded the duration of a task-irrelevant interval with much lower resolution than that of a task-relevant interval could account for RPEN responses and their relation to behavior [13].

Here, we investigate at a neural level whether RPENs implement an efficient code for reward value. We first derive the most efficient population of sigmoidally tuned neurons to encode rewards sampled from an arbitrary given distribution. We then apply this general framework to two data sets, one in mouse with a fixed reward distribution [15, 16] and one in monkey with a variable reward distribution [17]. We find that key properties of the efficient code are reflected in the data, suggesting that efficient coding could serve as a unifying principle. Finally, we develop learning rules for the efficient code.

We analytically derive the optimal population of neurons to encode rewards by extending the framework of Ganguli and Simoncelli [18]. Assuming that RPENs have a sigmoidal tuning curve for reward [Fig. 1A, 19, 20], we start with a large family of populations, within which we search for the most efficient one. We construct the family based on a base population of neurons on the unit interval. We then allow any smooth, monotone function to map from reward *R* to the unit interval and define the responses of the neurons in terms of the responses of the base population at the mapped location. Additionally, we allow an arbitrary scaling of the neurons’ response gains and of the density of neurons depending on their placement. In mathematical terms, we use a prototypical sigmoidal response function *h*_0_ on the unit interval to define the response of a neuron with midpoint at *μ* as:

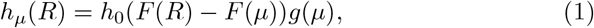

where 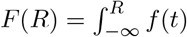 is the mapping from reward space to the unit interval. Its derivative *f* (*R*) ≥0 represents the conversion between reward space and the unit interval. *g*(*μ*) is the gain of a neuron with midpoint at *μ*. Finally, we define the density function *d*(*R*) as the probability that a neuron’s midpoint is placed at reward value *R*.

**Fig. 1.**
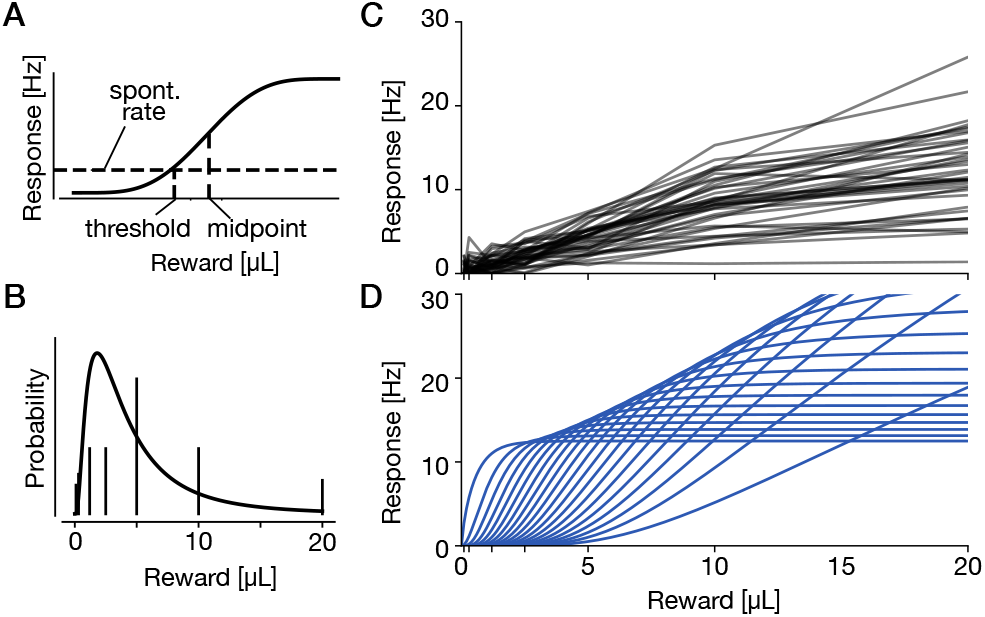
Experimental design and basic results of Eshel et al.[14]. A mouse was presented with variable rewards, while the responses of RPENs were recorded. **A**: Terminology for sigmoidal tuning curves as expected for reward prediction error neurons. The threshold is the reward value at which the spontaneous activity is reached. The midpoint is the reward value at which half the maximal response is reached. **B**: Rewards were drawn from a discrete distribution (vertical lines), which we approximate with a moment-matched lognormal distribution for our analyses (continuous line). **C**: Reward tuning curves of the 39 dopaminergic reward prediction error neurons measured by [14] after preprocessing of [15]. We additionally subtracted the minimal response across all trials for each neuron. **D**: Efficient code for the reward distribution in (A). For visual clarity, only 20 neurons are shown.

We then optimize the three functions to find the most efficient populations that can be generated this way to encode rewards from a distribution with density *p*(*R*) and cumulative density *P* (*R*) (see Online Methods). Efficiency here refers to the maximization of a measure of information given constraints on the expected total firing rate. This yields a family of equally efficient populations, which we parameterized by *α* ∈ [0, 1]:

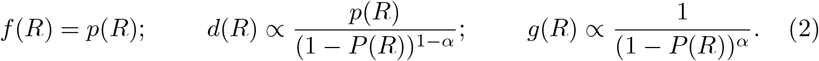

This solution extends the earlier result [18], which assumed a uniform density of neurons on the unit interval, effectively setting *d*(*R*) = *f* (*R*) in our formulation. Their solution is a special case of our family, with *α* = 1 and thus *d*(*R*) = *p*(*R*) and 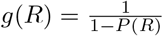.

### Application 1: Variable-reward task

We first compare the predictions of the efficient coding framework to data from the ‘variable-reward task’ of Eshel et al. [16] (Fig. 1B). In this task, mice were given one of seven reward magnitudes in each trial with certain probabilities, while RPENs responses were recorded. Fig. 1C shows the observed reward tuning curves of the RPENs. We derived the efficient code for a continuous approximation of the discrete probability distribution, in the form of a log-normal distribution matching the mean and variance of the discrete distribution (*μ* = 1.29 and *σ*^2^ = 0.71). The predictions for neural density and gain are shown in Extended Data Fig. 1. To compare our predictions to the data, we use the same number of neurons as in the dataset (*n* = 39) and adjust free parameters of the derived efficient population to match the measured population: We first set *α* to maximize the probability of the observed midpoints (whose distribution depends only on this parameter), resulting in *α* = 0.673. We define a neuron’s threshold as the point where the spontaneous firing rate is surpassed.

We fitted the spontaneous firing rate *r*^∗^ and *k*, a parameter controlling the slopes of all neurons, to the observed thresholds, resulting in the estimates 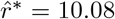 and 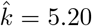. Finally, we set the constraint on the expected spike rate, *r*_max_, to match the average neural gain based on sigmoid fits to the tuning curves, resulting in 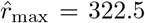 (see Online Methods for details). A subset of the resulting efficient code is shown in Fig. 1D. We now examine several properties of the efficient code and compare them to the data.

#### Prediction 1: RPEN midpoints cover the reward distribution

In the efficient code, RPEN midpoints are placed at specific quantiles of the distribution with a slight bias towards higher quantiles, yielding a distribution similar to the original reward distribution. Indeed, the midpoints of the observed neurons cover the range of the reward distribution, with a roughly similar distributional shape (Fig. 2A). The mean of the measured midpoints is 5.96, higher than the mean reward, which is 5.21 (*t*(38) = 0.848, *p* = 0.40). The fact that this difference is not significant is unsurprising, as for the measured of number of neurons, even the predicted midpoints are not significantly different from the mean reward (*t*(38) = 0.246, *p* = 0.81).

**Fig. 2.**
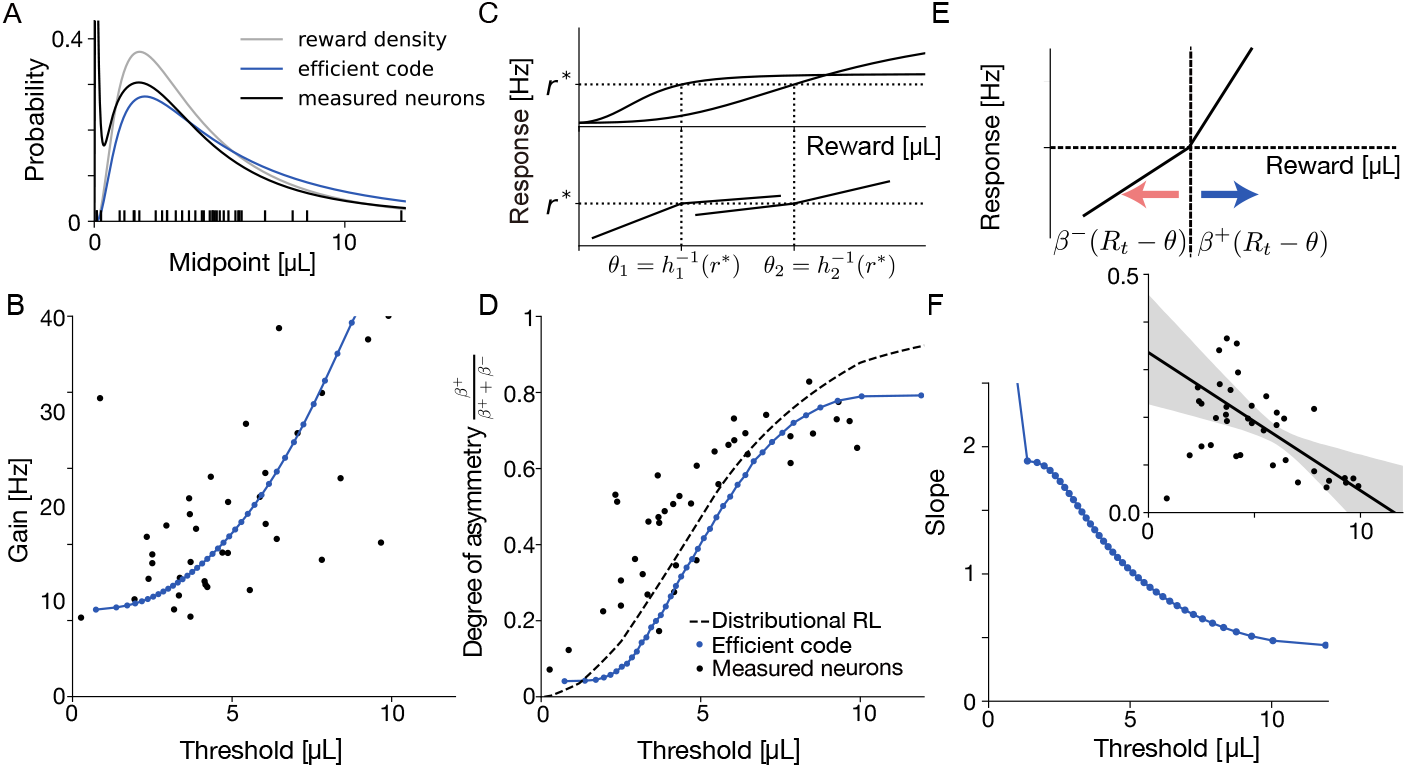
Comparisons between the measured neurons of Eshel et al. [16] and the efficient code. **A**: The distribution of the midpoints of the measured neurons covers the reward distribution with a slight upward shift (mean midpoint = 5.96, versus mean reward = 5.21, *t*(38) = 0.848, *p* = 0.402); the efficient code accounts for this (mean = 5.36). **B**: RPEN gain, i.e. maximal response of a fitted sigmoid function, plotted against threshold. There is a significant positive relationship (*r*(37) = 0.637, *p <* 0.001). The neurons of an efficient population (blue) again match the measured population (black) quite well. **C**: Efficient coding accounts for the relationship between asymmetry and threshold. The spontaneous activity *r*^∗^ is reached at *θ*_1_ within the concave part of the sigmoidal response function for a neuron with low threshold and gain, and at *θ*_2_ within the convex part of a sigmoidal response function for a neuron with high threshold and gain. **D**: Degree of asymmetry of neural responses 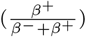 plotted against threshold. There is a strong positive relationship between these two variables (*r*(37) = 0.832, *p <* 0.001). The neurons of an efficient population (blue) closely approximate the the measured population (black). The dashed line is expected from ideal distributional TD learning. **E**: Distributional reinforcement learning accounts for the relationship between asymmetry and threshold: A reward value *R*_*t*_ shift the response function down proportional to *β*^−^ if below the threshold (*R*_*t*_ *< θ*) and up proportional to *β*^+^ if above the threshold (*R*_*t*_ ≥ *θ*). An equilibrium is achieved when the two expected shifts cancel each other. **F**: Slope of the sigmoid fit *a* plotted against threshold for the efficient code neurons. There is a significant negative linear relationship (*r*(37) = −0.915, *p <* 0.001). The inset shows the same plot for the measured neurons. There is a significant negative linear relationship in the data, too (*r*(37) = −0.550, *p <* 0.001).

#### Prediction 2: RPEN gain increases with threshold

In the efficient code, the gain is higher for neurons with higher reward thresholds (Fig. 1). The intuition is that those neurons respond to fewer rewards and can thus afford a higher gain with the same expected number of spikes. This effect does not occur for unimodal tuning functions [18, 21]. We observe the expected increase of gain in the data (Fig. 2B); to our knowledge, this empirical finding has not been reported before.

#### Prediction 3: RPEN tuning curve asymmetry flips with increasing threshold

In the efficient code, the increase of gain with threshold has implications for the shape of the RPEN tuning curves around threshold (Fig. 2C). As the gain increases, the spontaneous firing rate falls lower in the sigmoid shape. Thus, the threshold moves from the upper concave part of the sigmoid down into the convex part. In other words, neurons with low thresholds have concave tuning curves around their threshold and neurons with high thresholds have concave ones. To quantify the degree of such “asymmetry” of the neural responses, we followed previous work [15] and approximated the neural tuning curves with two linear response functions, one above and one below the spontaneous firing rate *r*^∗^ (Fig. 2C). This allows for an asymmetric response of the RPENs, with the slope above threshold *β*^+^ being different from the slope below threshold *β*^−^ [22, 23]. The ratio of slopes, ^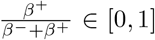^, can then be used as an index of asymmetry, which is close to 1 for concave tuning curves and close to 0 for convex tuning curves. Using this index, the efficient code predicts that asymmetry is higher for neurons with higher thresholds: the shape flips from convex to concave. This relationship is also observed in the data (black dots in Fig. 2D), as first reported by Dabney et al. [15].

The observed relationship can also be accounted for by the theory of distributional reinforcement learning (DRL; dashed line in Fig. 2D), which attributes it to an asymmetry of the update equations of the thresholds, such that neurons converge to different expectiles of the reward distribution (Fig. 2E). We will examine the relationship between efficient coding and DRL in detail below.

#### Prediction 4: RPEN slope decreases with threshold

In the efficient code, neurons with higher thresholds have a lower slope parameter of the sigmoid fit (Fig. 2F). This occurs because the predicted tuning functions are shallower in regions with lower reward density. In the measured neurons, we indeed find a significant negative correlation of slope and threshold (*r*(37) = −0.550, *p <* 0.001). To our knowledge, this empirical finding has also not been reported before. Overall, the measured neurons have shallower tuning curves than predicted by the efficient code. A possible explanation is that we did not take into account stochasticity in the RPEN inputs. Such stochasticity would produce random horizontal shifts of the sigmoid tuning curve, which, when averaged, would result in a lower slope.

### Application 2: Variable-distribution task

So far, we have examined RPEN responses to a fixed reward distribution. Rothenhoefer et al. [17] instead exposed two macaque monkeys to cues associated with rewards drawn from one of two distributions (either a “uniform” or a “normal” distribution) with the same mean (Fig. 3A-B). The main finding of Rothenhoefer et al. was that dopamine responses are amplified for rare rewards, producing steeper response functions for the “normal” distribution than for the “uniform” one (Fig. 3C-D). This suggests that RPENs encode the frequency of rewards.

**Fig. 3.**
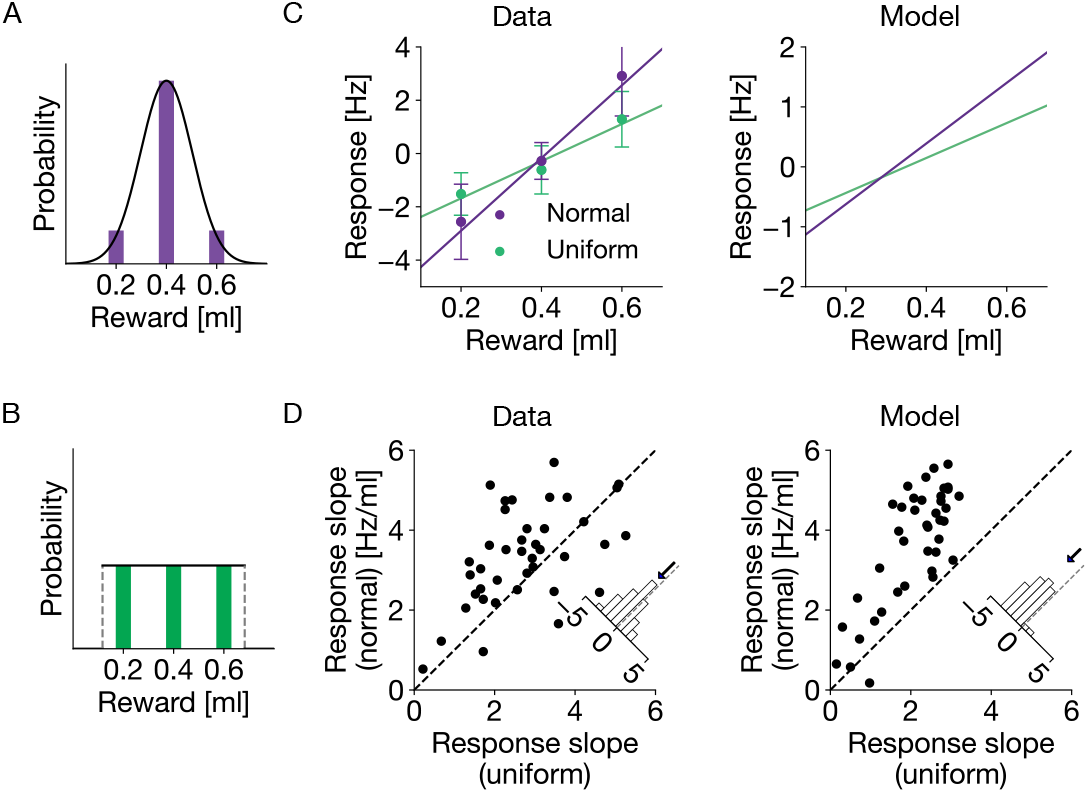
Efficient coding accounts for response slope characteristics in the Variable-distribution task. [17]. **A**: “Normal” distribution used in the experiment (purple) and moment-matched continuous distribution. **B**: Same for the “Uniform” distribution. **C**: Left: example neuron from Rothenhoefer et al.[17], showing steeper tuning for the “normal” than for the “uniform” distribution (left). Error bars represent ±1 s.e.m. across trials. Right: example neuron from the efficient code. **D**: Scatter plot of the response slopes of all neurons in the data [17] (left) and in the efficient code (right). Insets show the histogram of differences, and the downward arrow indicates the mean of it. The data showed a significantly different response slope with a mean difference of −0.605 (*t*(38) = −3.26, *p* = 0.002), and the model also demonstrated a significantly different response slope with a mean difference of −1.58 (*t*(38) = −10.4, *p <* 0.001).

The variable distributions make this data set an interesting additional test for the efficient coding framework. We derived an efficient code for a moment-matched continuous approximation of each distribution (Fig. 3A-B). We fitted the parameters *α, k*, and *r*_max_ to minimize the mean squared error between model and data on the response slopes for the individual distributions. The resulting estimates were 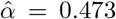,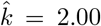 and 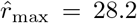 (Extended Data Fig. 2). As in Application 1, we find that the midpoints of the neurons in the efficient code cover the range of the reward distribution (Extended Data Fig.2B and D). Moreover, the efficient coding model successfully reproduced the main finding (Fig. 3C-D): a majority of neurons in the efficient code (39 out of 40) exhibited steeper slopes for the “normal” distribution, in agreement with the data (31 out of 40).

### Learning rules for the efficient code

Distributional RL [15] proposes that RPENs update their thresholds in proportion to the difference between the received reward and their thresholds, with different proportionality factors for positive and negative differences (Fig. 2E). This learning rule leads those thresholds to converge to the expectiles of the reward distribution. This is very similar to the prediction of the efficient code, which places neuron’s midpoints at specific quantiles of the distribution. With the additional assumption that the slope of the reward tuning curve above and below threshold is proportional to the learning rate, distributional RL accounts for the observed relationship between threshold and asymmetry. However, without modification, it does not account for the relationship between gain and threshold or the relationship between slope and threshold. Since the efficient code does account for these observations, a good way to consolidate the two theories is to extend the DRL learning rule such that the gain and slope of neurons also converge to the parameters for the efficient code.

To learn the placement of neurons on the reward axis, we can use the asymmetric DRL learning rules. To converge to the quantiles of the distribution, we shift the neuron to higher reward values by a step size *b* and towards lower reward values by a different step size *a* if a reward below the midpoint occurs (Fig. 4 A). If we set *a* and *b* such that *ap* = *b*(1 −*p*), this procedure converges to the *p*th quantile of the distribution. In the DRL formulation, neurons instead update their position proportional to the difference between their location and the presented reward and thus converge to the expectiles of the distribution. This changes the population only slightly, but achieves similar efficiency as the populations placed at the quantiles (Extended Data Fig. 3). It should also be kept in mind that both the step function that we apply for the quantiles and the linear weighting functions that yield expectiles are idealizations.

**Fig. 4.**
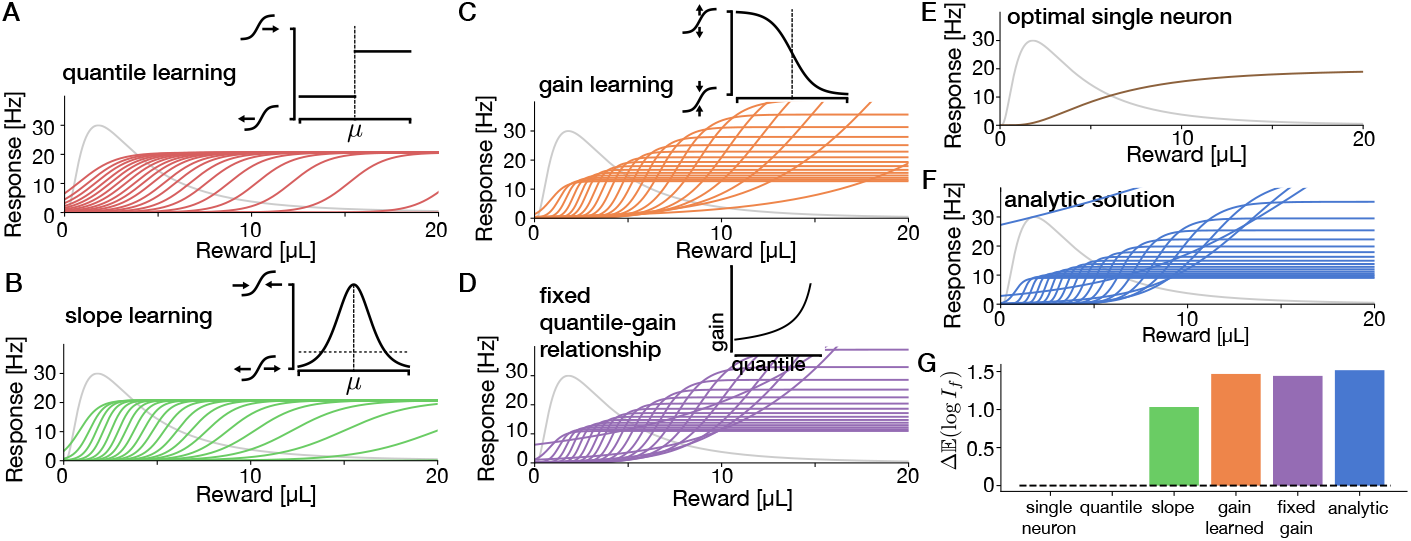
A combination of learning rules to learn the efficient code. Each panel shows the steady state distribution population of 20 neurons after learning based on 20 000 reward presentations with an inset illustrating the learning rule. **A**: Learning the position on the reward axis for the neurons to converge to the quantiles of the distribution. This learning rule is similar to the distribution RL learning rule. **B**: Additionally learning the slope of the neurons to be proportional to the local density by increasing the slope when the reward falls within the dynamic range and decreasing otherwise. **C**: First method to set the gain: iterative adjustment to converge to a fixed average firing rate. **D**: Second method to set the gain: Use a fixed gain per neuron based on the quantile it will eventually converge to. **E**: The efficient tuning curve for a single neuron. **F**: The analytically derived optimal solution. **G**: Comparison of the different populations in the overall information transfer with the same number of neurons (20) and expected firing rate (8.27 per neuron as in the fit to the measured data).

In the efficient code, RPEN slope should be proportional to the local probability density of rewards. To learn the slope, we can estimate the local probability density based on the probability that a reward within the dynamic range of the neuron occurs, i.e. in the range where the neuron has a high slope. As the dynamic range is inversely proportional to the slope, it should be inversely proportional to the local reward density as well, which means that the probability of a reward in the dynamic range should be constant. A simple learning rule to achieve this is to increase the slope and shrink the dynamic range when a reward falls into the dynamic range and decrease the slope and increase the dynamic range when a reward falls outside the dynamic range. When we allow a gradual definition for falling into the dynamic range, a natural choice is to use the derivative of the sigmoid function; this is the kernel we chose for our simulations (Fig. 4B). Any other kernel around the midpoint of the sigmoid whose width scales inversely with the slope would also work.

Finally, we need to adjust the gain of the neurons to match the efficient coding solution. For this purpose, we present two solutions that work similarly well but that are based on different biological mechanisms. In the first method, we learn an estimate of the expected firing rate and set the gain based on this estimate. Effectively, this results in the inverse of the sigmoid shape for updating the gain with high rewards reducing a neuron’s gain and low rewards increasing its gain (Fig. 4C). In the second method, we couple the fixed gain to the quantile the specific neuron should converge to based on its learning asymmetry (Fig. 4D).

These learning rules for midpoints, slope, and gain together produce a population of neurons similar to the analytical efficient coding solution (Fig 4C, D, F), with similar efficiency (Fig. 4G). These populations clearly beat the best possible solution that assumes the same tuning function for every neuron (Fig. 4E,G). Thus, these rules successfully extend the DRL learning rule to produce an efficient code.

## Discussion

We have presented evidence that RPENs implement an efficient code for reward. Starting from a normative account of the tuning of RPEN responses, we were able to account for five empirical findings across two tasks. Thus, efficient coding could serve as a unifying principle underlying the response characteristics of RPENs.

Our results are robust to changing the measure of information used for optimizing the population, the assumed distribution of neural responses, or the assumption of a sigmoid shape. Different objective functions or different response distributions lead to different relationships between the density of the reward distribution and the optimal density and gain of neurons [18]. However, neural density always increases with reward density, as it is more efficient to focus neural resources on probable rewards. Analogously, all objectives lead to higher gains at higher thresholds, as gain increases are “cheaper” for neurons with higher thresholds. Furthermore, changing the mathematical form of the shape of the response function does not change the density and gain predictions, as those were derived independently of the shape, and only leads to slightly different predictions for the asymmetry of the responses around the threshold. For any shape that is convex in its lower response range and concave in its upper response range, the qualitative argument for the dependence between threshold and asymmetry will hold.

Our work has limitations. To simplify our analysis, we restricted ourselves to a task with a single reward supplied in each independent episode without any action required by the animal. In this simple case, the reward received and the change in value of the current state are confounded. Thus, all our analyses are agnostic to this distinction. In multi-step [24] and delayed-reward tasks [25], RPENs take future rewards into account with temporal discounting. The efficient coding framework we presented here would need to be modified for a temporally discounted value code. Moreover, our theory covers only the phasic reward prediction error response of RPENs. The ramping activity of RPENs [e.g. 26, 27] will require additional explanation, most likely including an encoding of reward timing [28]. In fact, it has been proposed that RPENs encode elapsed time more generally [29], and that this encoding is tuned to task properties in a manner reminiscent of efficient coding [13].

The efficient coding hypothesis is compatible with reinforcement learning. Specifically, we extended the learning rule from DRL [15] to account not only for the relationship between asymmetry and threshold but also for the three other tuning properties predicted by the efficient code. Besides DRL, there are two alternative explanations of the data by Eshel et al. [16]: the Laplace code [30] and normalized reinforcement learning [31]. In the Laplace code, TD-learning neurons with different parameters are used to encode the timing and a whole distribution of rewards in the future by representing an analogue of the Laplace transform in the neural responses. Individual neurons still have a sigmoid response curve in this code and the dependence between asymmetry and threshold can be created similarly as in the efficient code by cutting the curve at different response levels. In their formulation, they subtract the expected response from each neuron instead of changing the gain (as we predict and observe). Normalized reinforcement learning [31] proposes that RPENs perform divisive normalization with different half saturation constants. This yields sigmoid (Naka-Rushton [32]) neurons with different thresholds, and the asymmetry around those thresholds is explained again by cutting the sigmoids at different heights. Neither of these explanations explicitly examines code efficiency or attempts to account for the dependencies of gain or slope on threshold.

Future work could examine how learning rules that produce an efficient code can be implemented in biologically realistic circuits (perhaps similar to [33–36]), empirically distinguish between candidate learning rules, and examine how RPENs can relatively quickly switch from one reward distribution to another.

## Supplementary information

This article is accompanied by Supplementary Information.

## Acknowledgments

We thank Hsin-Hung Li for valuable discussions.

## Declarations

### Availability of data and materials

No new data was measured for this project. The data collected by Eshel et al. that we analyze here are kindly made available by Dabney et al. [15] at https://doi.org/10.17605/OSF.IO/UX5RG.

### Code availability

The code to recreate our analyses is available at https://github.com/dongjae-kim/efficient-coding-dist-rl

### Author contributions

HS derived the efficient code; HS and DK analyzed the neural data. WJM supervised this project. All authors wrote the manuscript.

## Online Methods

### Analysis of neural data

#### Variable-reward task

We used the data from the variable-reward task by [16] as preprocessed by [15]. For details of data acquisition, please refer to [16]. Five mice were subjected to a variable-reward paradigm incorporating two experimental conditions: a no-odor condition and odor condition. During each trial, one of seven reward magnitudes (0.1, 0.3, 1.2, 2.5, 5, 10, and 20 *μ*L) was randomly chosen with probabilities of 0.066, 0.091, 0.15, 0.15, 0.31, 0.15, and 0.077 respectively. Electrophysiological data were recorded from *n* = 40 ventral tegmental area dopaminergic RPENs. From each of the mice, 3, 6, 9, 16, and 6 neurons were selected respectively. One neuron was excluded because it had never shown a single response larger than its baseline for any reward magnitude. We used the data as preprocessed by Dabney et al.[15]. As part of this preprocessing, on each trial, a pre-stimulus response was subtracted from the later responses. As this procedure yielded negative firing rates, we additionally subtracted for each neuron the overall smallest response from all its responses.

#### Variable-distribution task

To further investigate response of dopaminergic RPENs with different reward distributions, we analyzed the data from the variable-distribution task presented in Rothenhoefer et al. [17]. For detailed data acquisition methods, we refer to that paper. Two macaque monkeys were subjected to a variable-reward paradigm, where cues indicated rewards from either a uniform or a normal distribution. In each trial, rewards of 0.2, 0.4, or 0.6 mL were delivered based on a uniform distribution (1/3 chance for each amount) or a normal distribution where 0.2 and 0.6 mL had a 2/15 probability and the 0.4 mL reward had an 11/15 probability. Electrophysiological data from RPENs were collected to monitor dopamine responses. Peri-stimulus response was subtracted from the later responses, and smoothed as in the variable-reward task.

### Logistic fits

For each neuron separately, we fitted logistic sigmoid functions 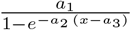 to the responses. We denote the gain *a*_1_, the slope *a*_2_, and the midpoint *a*_3_ collectively by **a**. To estimate the parameters, we used maximum-likelihood estimation based on Poisson noise, so that

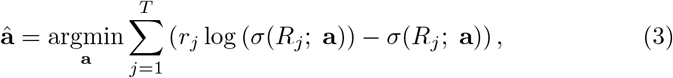

where *r*_*j*_ is the measured firing rate of a neuron responding to the reward magnitude *R*_*j*_ on the *j*th trial. We proceeded to compare the fitted parameters between the data and the efficient code.

### Fitting asymmetric slopes

To enable comparisons to the asymmetry data of [15], we repeated their analysis and applied the same asymmetry calculation to the simulated neurons in our efficient code.

First, we subtracted a baseline from the smoothed PSTHs for each neuron and each trial. The baseline was the mean firing rate over the 1000 ms before stimulus onset. We then estimated asymmetric slopes, *β*^+^ and *β*^−^ by separately fitting linear functions to the responses above and below threshold. Following Dabney et al. [15], we used estimated utility space as x-axis for this step, but the results are not sensitive to the choice of utility function.

In math terms, the slope above threshold *β*^+^ and the slope below threshold *β*^−^ for responses *y*_*j*_ to rewards *R*_*j*_ are thus defined as:

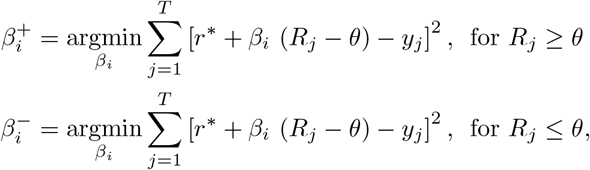

where *r*^∗^ is the spontaneous firing rate and the sum over *j* runs over the *T* trials.

To achieve more efficient fitting than Dabney et al. [15], we derive analytical solutions for *β*^+^ and *β*^−^ for a given *θ*, which are:

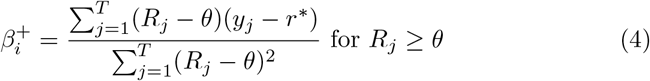

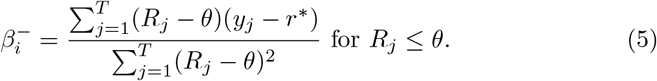

Thresholds – called reversal points in [15] – were then chosen to minimize the overall sum of squares, by sampling 20,000 thresholds from an uniform distribution between the range of rewards and choosing the best-fitting one. Dabney et al. used the same random sampling procedure for *β*^+^ and *β*^−^.

For fitting our simulated neurons, we set the threshold 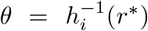 and *y*_*j*_ = *h*_*i*_(*v*_*j*_) and fit the *β*s based on the results for 10^5^ trials each above and below threshold, sampled from the approximated log-normal reward distribution restricted to the range above and below threshold respectively.

### Derivation of the efficient code

To derive the efficient code for reward, we extend the framework for analytical solutions proposed by Ganguli and Simoncelli [18]. In this framework, a uniform standard population is defined on the unit interval [0, 1] ^1^. By using an adaptable mapping between the unit interval and the stimulus space and functions that transform the population, we can define a broad class of populations among which we can find the most efficient ones analytically. We extend this framework by allowing the distribution of neurons to be non-uniform, which decouples the local slope and density of neurons that are coupled in the original framework.

### Definitions

We start by defining a “standard” population of RPENs. This standard population is a collection of sigmoidal tuning curves *s*_*ω*_ : [0, 1] → [0, 1] indexed by their ‘center’ *ω ∈* [0, 1]. Individual neurons are assumed to be monotonically increasing functions whose derivatives are unimodal functions with their peak near *ω*. Additionally, we assume that the overall Fisher information *I*_F_(*x*) of this population about the position *x∈* [0, 1] and the overall increase in firing rate are independent of *x* i.e.:

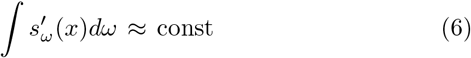

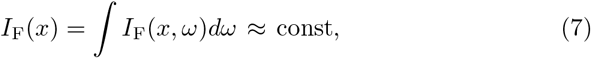

where *I*_F_(*x, ω*) is the Fisher information provided by the tuning function with center *ω* and 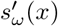 is the derivative of the tuning function.

Finally, we assume that the total amount of Fisher information provided by each neuron is approximately constant:

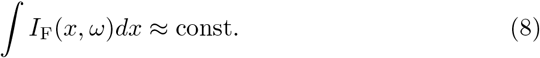

One simple way to construct a population that fullfills these criteria is to start with a prototypical sigmoid function *s*_0_ centered on 0 and defining the other tuning curves as *s*_*ω*_(*x*) = *s*_0_(*x* −*ω*). This construction often leads to strong edge effects at 0 and 1, because a substantial part of the sigmoid response curves near the borders will fall outside the [0, 1] interval. We ameliorate these effects by using a more sophisticated population based on cumulative beta distributions below. Nonetheless, this simple construction illustrates the intuition behind a uniform population of sigmoidal neurons well and was actually used by Ganguli and Simoncelli in their implementation.

To map between the unit interval and stimulus space we use a strictly monotonic function 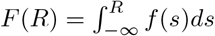 from the stimulus space into the unit interval, parameterized by its derivative *f* (*s*) *>* 0, which we will optimize below. As *F* is strictly monotonic, it is also invertible, with its inverse *F* ^−1^ mapping the unit interval into stimulus space. Additionally, we allow a gain function *g >* 0 of stimulus space, which scales the tuning curve of neurons placed at different positions in stimulus space.

With these definitions, we can now define the tuning function *h*_*μ*_ of a neuron with midpoint at stimulus level *μ*:

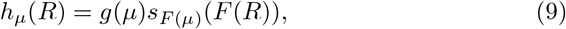

i.e. we map both the positions of the neurons and the stimuli into the unit interval, evaluate the sigmoids there and scale the result with the gain.

Up to this point our definitions are equivalent to those of Ganguli and Simoncelli [18], who now assume that the neurons are equally spaced in the interval. This additional assumption makes it impossible to place more neurons in some stimulus range without changing their shape. This removes one of the possible adaptations of a neural populations mentioned by other authors like Wei and Stocker [37] for example. To allow different distributions of the neurons, we instead assume that the *μ*_*i*_ are drawn from a distribution over reward space with density *d*, which we will optimize below. The equal placement assumed by Ganguli and Simoncelli is equivalent to the assumption *d*(*R*) = *f* (*R*), which yields a uniform distribution over the placements in the unit interval *ω*.

### Optimization Objective

We further assume that the firing rates of the RPENs are subject to independent Poisson noise, which is the simplest noise assumption. We can then calculate the Fisher information *I*_F_(*R*; *μ*) provided by a neuron centered on *μ*:

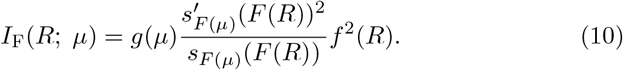

We then optimize the expected value (over both *R* and *μ*) of the logarithm of the Fisher information *I*_F_, based on *N* neurons:

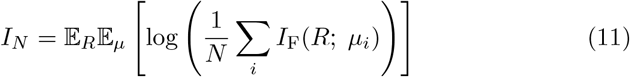

This objective was originally proposed as a lower bound on the mutual information between the neural responses and the stimulus [18, 38, 39], which becomes tight for small Gaussian shaped errors on the encoded value. However, some later publications cast doubt on this interpretation: In particular, [40] interprets this equation as an *upper* bound on mutual information instead. Additionally, low firing rates and/or steep changes in firing rate can lead to this approximation being highly inaccurate, and this realm is actually reached for cases in which neurons are measured only for short times [41]. Nonetheless, *I*_*N*_ remains the best Fisher information-based approximation to mutual information, and we follow Ganguli and Simoncelli [18] in optimizing it. They also tried a few other functionals of Fisher information and got qualitatively similar results. Mutual information itself is unfortunately untractable for these derivations.

Assuming further that there is a large population of RPENs, we can now look at this distribution in the limit of infinitely many neurons. According to the strong law of large numbers, the logarithm of the mean in Eq. (11) almost surely converges to the logarithm of the expected Fisher information, because the Fisher information provided by each single neuron has finite variance:

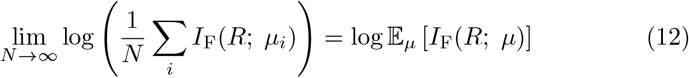

Because this is a constant with respect to *μ*, we can drop the outer expected value, yielding:

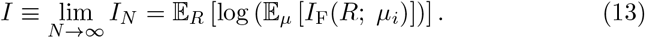

To make this expression amenable to optimization, we simplify it as follows:

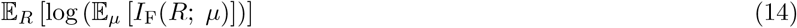

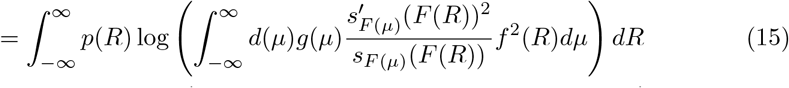

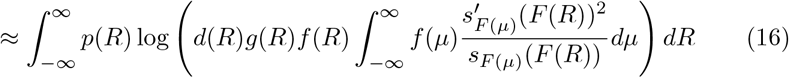

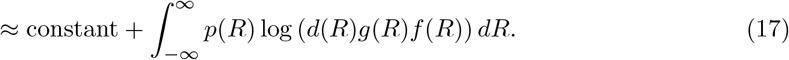

Here, we eliminated the inner integral through two steps, which are only approximately correct: First, we observe for sigmoidal functions, that 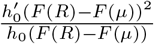 is a unimodal function that is centered around *R*, which effectively acts as a smoothing kernel that is convolved with the rest of the integral. If *d*(*μ*), *f* (*μ*) and *g*(*μ*) change little over the range of this smoothing kernel, we can replace them by their values at *R* and pull them out of the integral. Second, the integral over *μ* we are left with is (approximately) shift-invariant, i.e. the same for all *R* and invariant to the monotonic transformation *F*, if the original population is (approximately) uniform as defined in Eqs. (7) and (8).

The final optimization problem is now to optimize this approximation of *I* with respect to the three functions *d, g*, and *f* :

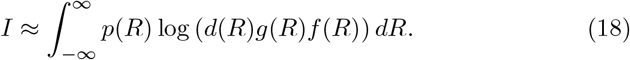

Additionally, we enforce that the expected number of spikes E[*r*] under the prior distribution is smaller or equal to a bound *r*_max_. The expected number of spikes is:

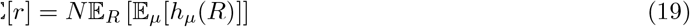

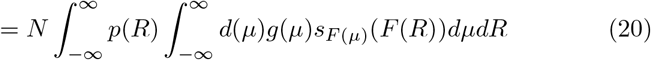

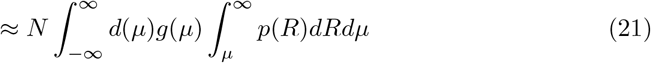

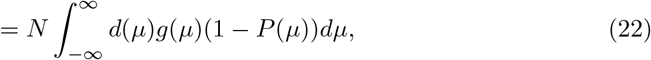

where *P* is the cumulative density function of the reward distribution. In Eq. (21), we approximated the sigmoid with a step function at *μ*. This approximation is close if *p*(*R*) is near constant over the range where the sigmoid *h*_*μ*_ changes from 0 to 1. This is the case if the sigmoids are reasonably steep.

There are two further constraints, one to make *d* a valid probability density and one to limit the range of *F* to the unit interval:

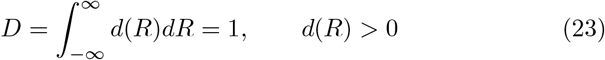

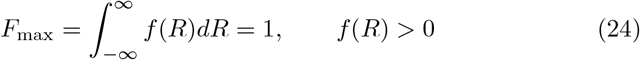

### Solving the Optimization

Now we are ready to solve this optimization problem using Lagrange multipliers. To do so, we first compute the functional derivatives of our objective function:

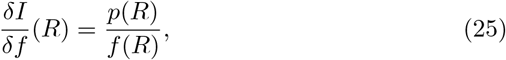

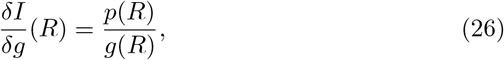

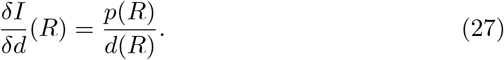

Similarly, we compute the functional derivatives of the constraints. First, for the expected spike rate:

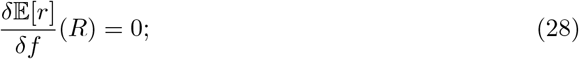

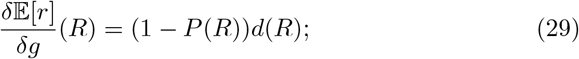

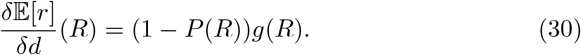

Second, for the integral of the density *D*:

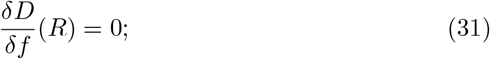

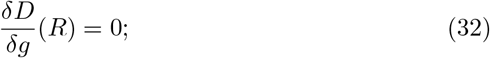

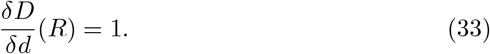

And finally, for the upper end of the interval reached by *F, F*_max_:

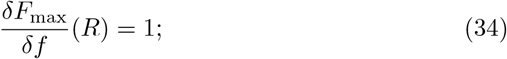

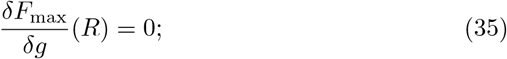

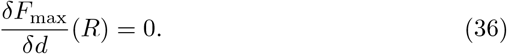

With these derivatives in place, we solve the optimization problem using Lagrange multipliers *λ*_*R*_, *λ*_*D*_, *λ*_*F*_, which results in the following three equations:

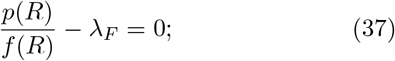

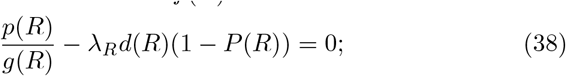

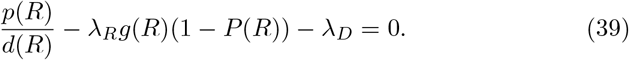

From Eq. (37), we obtain

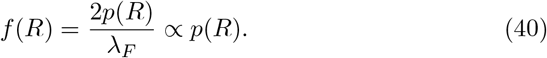

From Eq. (38), we obtain

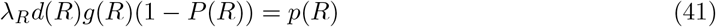

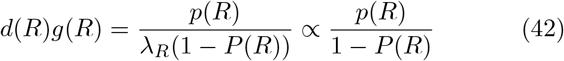

By inserting this result into Eq. (39), we see that it is guaranteed with *λ*_*D*_ = 0.

We conclude that under a constraint on the expected number of spikes, the neural population that optimizes the Fisher information bound on mutual information has the following properties:

- The inverse slope of the sigmoids is proportional to the probability density.
- The product of the probability density for neurons with a given midpoint and the gain for neurons with this midpoint is equal to the ratio of the reward probability density and the probability that rewards larger than the midpoint appear.

The formulation by Ganguli & Simoncelli [18] with equally spaced neurons in the unit interval enforces the additional constraint that *d*(*R*) ∝ *f* (*R*) and thus arrives at the special case of *d*(*R*) ∝ *p*(*R*) and *g*(*R*) ∝ (1 − *P* (*R*))^−1^.

### Splitting density and gain

In our solution, *d* and *g* entirely compensate for each other. Due to the fact that their product needs to be proportional to a product of the two factors *p*(*R*) and 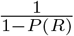, a natural choice to parameterize a family of solutions is to use a parameter *α* ∈ [0, 1] that trades off the distribution of the two factors to density and gain:

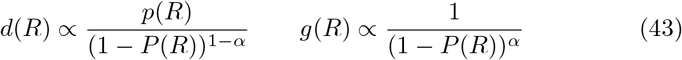

Effectively, *α* trades off the gain increase at higher thresholds against placing more neurons at higher thresholds. We illustrate this for several different reward distributions and values of *α* in Fig. 5.

**Fig. 5.**
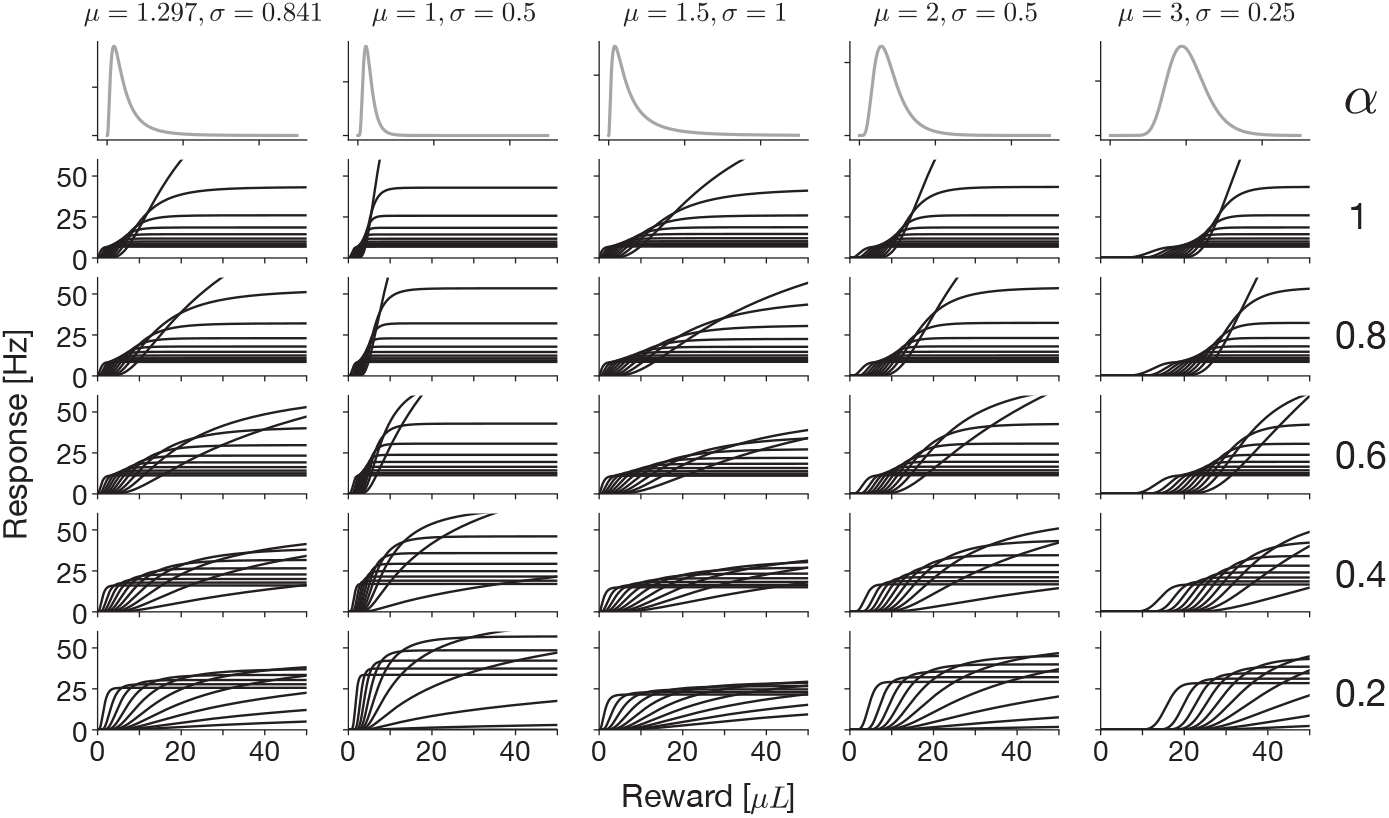
Illustration of the solution of the efficient coding problem, varying *α* (rows) and the reward distribution (columns). The reward distributions are all log-normal distributions with their pdfs and parameters plotted at the top.

Other ways of writing 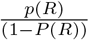 as a product of neural density and gain are possible, but the split we chose has two convenient properties. First, the density *p*(*R*) appears only as a whole factor, which corresponds to the density of the reward distribution actually observed by the animal. This allows realistic learning rules for the neurons, as we derive below. Second, this formula for the density yields a valid density when *α >* 0 independent of the reward distribution, so that the same split can be applied to all distributions. We can derive this fact by looking at the uniform distribution on [0, 1]. For this reward distribution, the density of neurons becomes (1 −*x*)^*α*−1^, whose integral over [0, 1] is:

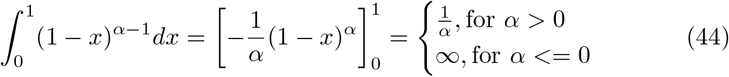

Thus, the valid range for *α* for the uniform distribution is exactly the positive numbers. For any other reward distribution, we can transform the distribution of neurons for the uniform distribution with the inverse cumulative density function *P* ^−1^. This yields samples with the desired density 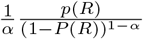, which must therefore be a valid density.

### Choosing a Solution

We fitted *α* using only the midpoints of the measured neurons. To do so, we extracted the midpoints of the measured from logistic function fits as described above. We then estimated *α* using maximum-likelihood estimation, resulting in 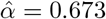.

The requirements we put forth in Eqs. (6)-(8) are approximately met by many populations of sigmoid-shaped tuning curves. Thus, the exact shape and in particular the slope of the sigmoid are not fixed by these efficient coding considerations. For all optimal populations in this paper we use the cumulative density functions of beta distributions, because these sigmoid functions are defined on the unit interval [0, 1] and become exactly 0 and 1 at the ends of the interval, which yields good boundary behavior. These sigmoid functions are not exactly shifted versions of each other as we assumed in the derivations, but functions that are shifted versions of each other lead to relatively strong boundary effects that produce worse deviations from the theory. To place a neuron at a specific position *p* ∈ [0, 1], we set the parameters such that *p* is the mean of the beta distribution (*a* = *kp, b* = *k*(1 −*p*)). Thus, we are left with one more parameter *k >* 0, which controls the slope of the sigmoid functions. As the efficient coding scheme does not determine what the overall slope should be, we set *k* = 5.20 to fit the threshold distribution of measured neurons as described below. We do not expect other shapes to behave fundamentally differently.

We used a fixed spontaneous firing rate *r*^∗^ for all RPENs. We fitted *r*^∗^ and the slope parameter *k* to minimize the mean squared error between the thresholds of the efficient code, 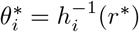), and the thresholds of measured neurons:

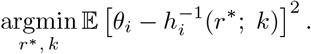

The resulting estimates were 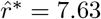 and 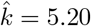.

Finally, the efficient code has a parameter *r*_max_ that restricts the budget for spikes. In the efficient code, the gain of all neurons is proportional to this parameter. To fit this parameter, we fit sigmoids to both the empirical and the efficient tuning curves as described above and added the squared error of the neuron-averaged gains to the optimization objective.

### Checking the solution

We performed several checks to confirm that the derived population is indeed efficient given the reward distribution. To do so, we compared the amount of information communicated to some other populations with the same number of neurons and the same expected number of spikes (Fig. 6). The optimized populations retain more information than other populations. The population with the optimized *α* = 0.673 is equally efficient as the solution of Ganguli and Simoncelli [18], which corresponds to *α* = 1.

**Fig. 6.**
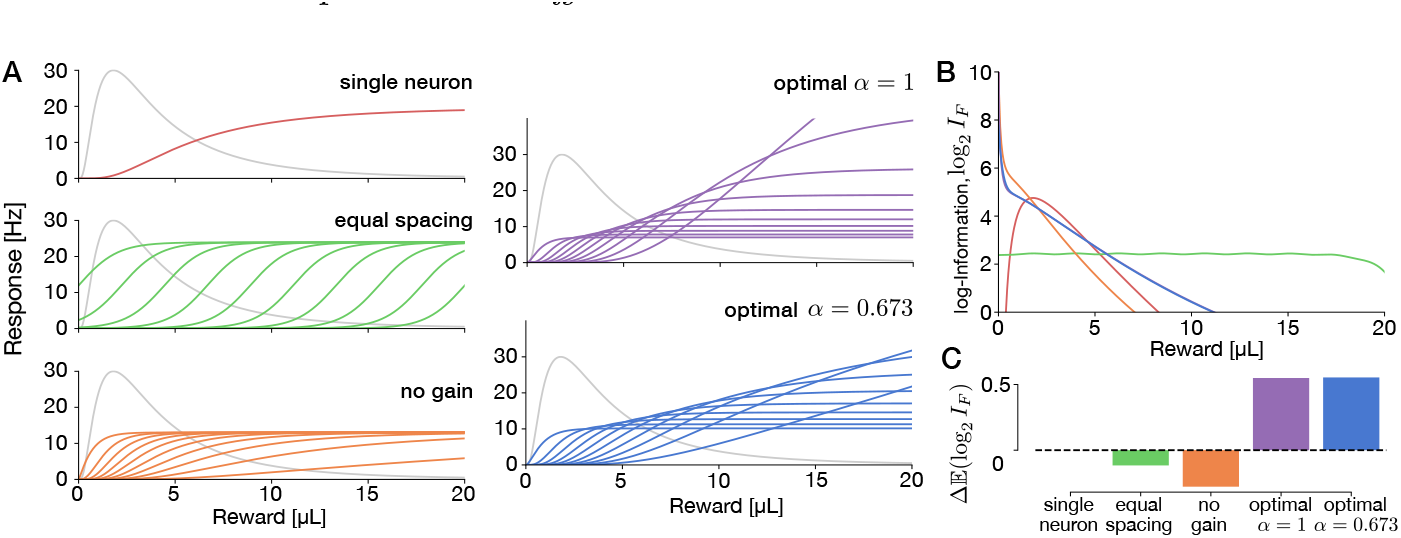
Comparing encoding populations for reward with 10 neurons and the same expected number of spikes. **A**: Compared neuronal populations: **single neuron: All neurons share the same response curve, optimized to maximize transferred information. equal spacing: neurons tile the space, not optimized.no gain: positions and slopes are optimized, but all neurons have equal gain. optimal** *α* = 1**: fully optimized population as derived previously[18] with density proportional to the distribution. optimal** *α* = 0.673**: Equally optimal distribution but with** *α* **fit to match the midpoint distribution for the optimal code and the experimental data. B: Fisher information as a function of reward for each of the populations. C: Expected logarithm of Fisher information under the reward distribution relative to the single-neuron case**.

### Other Objectives

Our predictions stay qualitatively similar for optimization objectives that transform the Fisher information in different ways, such as the discrimax objective studied by [18]. In particular, our results can be derived whenever the loss is an expected value over a convex function of Fisher information, which goes to infinity at 0 and is monotonically decreasing. Then there are diminishing returns for increasing the fisher information at any specific reward value, but we need to achieve some nonzero Fisher information for finite loss. Then the optimal population will always be a trade-off between distributing the Fisher information broadly and the lower cost for higher thresholds.

As the Fisher information itself and the expected spike rate both depend on the density and gain only through the product of density and gain, we will find a trade-off between density and gain for any objective function defined in this way.

### Learning rules for the efficient code

Here, we define the learning rules for the efficient code exactly and show that they converge to the desired population.

#### Midpoints

We start with a learning rule that makes the midpoints converge to the quantiles of the reward distribution, so that their density is proportional to *p*(*R*). This covers the *α* = 1 case of Eq. (2). Then, we will describe how to achieve other *α*-values by choosing the quantiles from a specific non-uniform distribution.

To create a neuron that converges to the quantile *q* of the reward distribution, we can adjust its midpoint *μ* in analogy to the distributional RL learning rule: When a reward above *μ* occurs, we increase *μ* by a constant value *γ*_+_, if a reward below *μ* occurs, we decrease *μ* by a (usually) different value *γ*_−_:

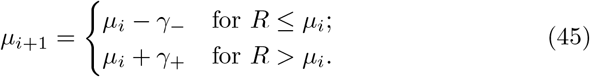

The expected change of the midpoint with this learning rule is:

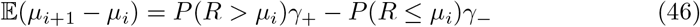

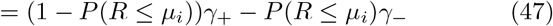

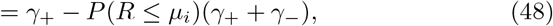

which is 0 exactly if the current quantile of *μ, P* (*R* ≤ *μ*_*i*_), satisfies

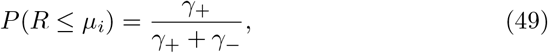

negative if it is larger, and positive if it is smaller. If we set *γ*_+_ and *γ*− such that 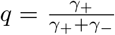, the neuron with these parameter values will converge to (or oscillate around) the desired quantile. By setting the distribution of *γ*_+_ and *γ*_−_ values across the population of neurons, we can thus enable convergence to an arbitrary distribution of quantiles. Choosing a uniform distribution solves the *α* = 1 case.

To enable convergence to a distribution with density proportional to 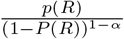 for *α* ≠1, we can choose quantiles proportional to 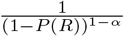. By the transformation theorem for probability densities, this will result in a distribution with the desired density.

#### Slopes

The slopes need to converge to values proportional to the local density of rewards, see (40). As in kernel density estimation, an estimate of the density around the midpoint of a neuron *μ* can be found by evaluating a positive kernel function *K* with integral 1 at the observed reward values *R*_*i*_:

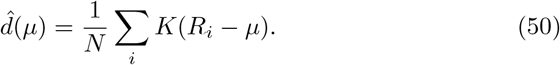

From this equation, we could already derive a simple learning rule for the slope of the sigmoids, namely setting them based on a running average of these kernel evaluations. This would imply a fixed kernel size for any reward distribution though, which would be suboptimal if the variance of reward distributions varied. To avoid this problem, we can use learning rule that adapts the kernel size simultaneously with the slope. We set the width of the kernel *h* anti-proportional to the slope *b* of the sigmoid function. Thus, the tuning function and the kernel are stretched and shrunk equally, such that *bh* = constant. Then we get:

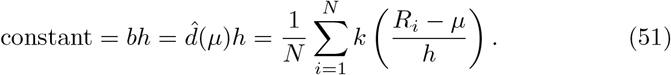

In other words, we need to define a learning rule, such that 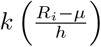 converges to a constant. This can be achieved for any constant *c* ∈ [0, *k*(0)] by changing the slope and kernel width proportional to 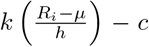. As *h* and *b* need to stay positive, we perform these updates in log-space, i.e. multiply both by 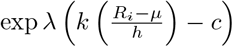 for some small learning rate *λ*.

#### Gain learning

As the first of two ways of adjusting the gain of a neuron, we can use the expected firing rate before scaling with the gain. This can achieve convergence to the efficient code, because the factor 1 −*P* (*μ*) arises as an approximation of the expected firing rate of a neuron in Eq. (22) before multiplication with the gain. To learn this value, a neuron can compute a running average of the unscaled neural responses 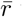 with some learning rate *λ*_*r*_:

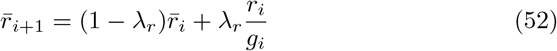

Plugging this into the efficient code solution yields:

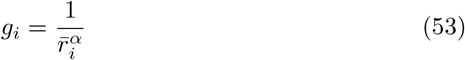

With *α* = 1, this learning rule would yield the same expected firing rate for all neurons. With *α <* 1, it yields a less extreme growth in gain.

#### Gain coupled to quantile

Another way of adjusting the gain is by setting a fixed gain based on the quantile a neuron is meant to converge to. As the quantile fixes *P* (*μ*), we can read off the correct value from the analytical efficient coding solution, i.e. set the gain *g* equal to 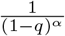 for neurons that are meant to converge to the quantile *q*.

### Expectiles instead of quantiles

For the figures in the main text, we placed neurons’ midpoints at the quantiles of the reward distribution as predicted by maximizing coding efficiency. As distributional RL predicts that neurons should converge to expectiles instead, we tested the efficiency of such a code in Extended Data Fig. 3. The efficiency of the code is slightly lower for the expectile-based code, due to a slight shift of midpoints towards the mean of the distribution. These differences are small and the code does not show qualitative differences. We understand them as essentially equivalent.

## Extended data figures

**Extended Data Fig. 1.**
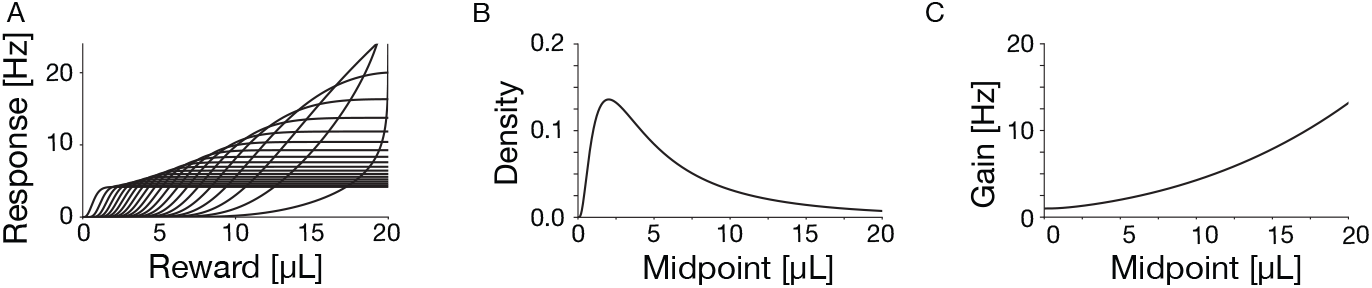
Efficient code for the variable-reward task [16]. **A**: Tuning curves. For clarity, only 20 of 39 neurons are shown. **B**: Density of neurons as a function of midpoint. **C**: Gain as a function of midpoint.

**Extended Data Fig. 2.**
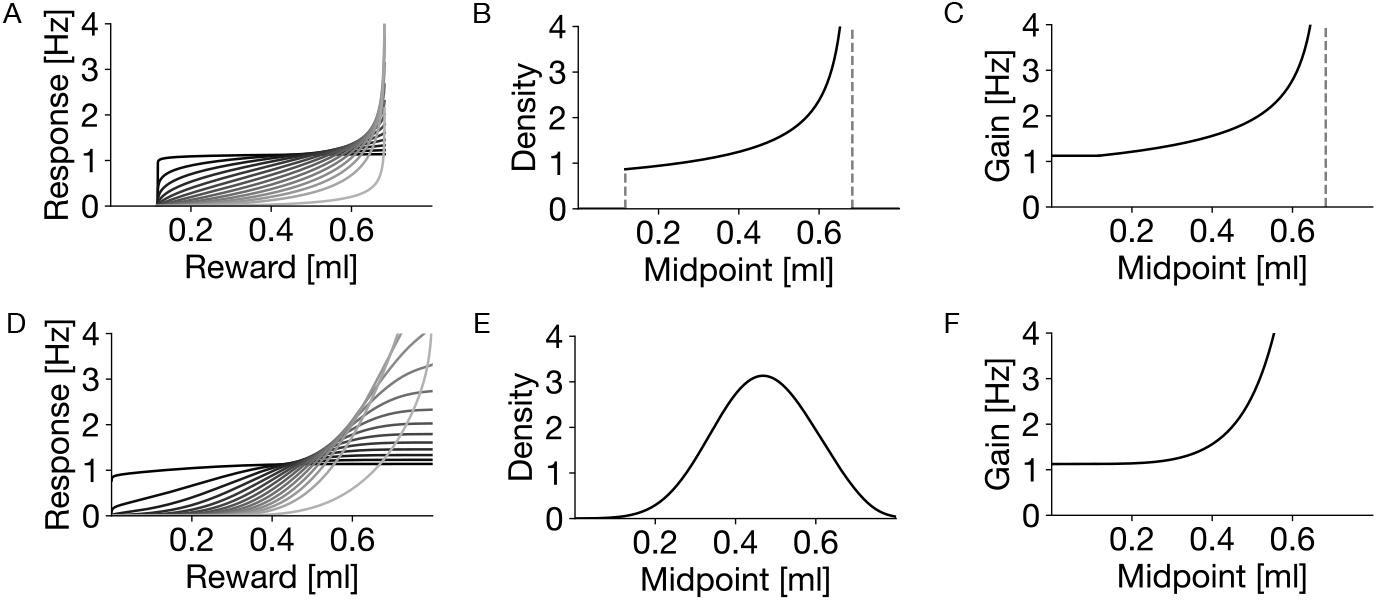
Efficient code for the variable-magnitude task [17]. **A-C**: Efficient code for the uniform distribution. **D-F**: Efficient code for the normal distribution. **A**,**D**: Tuning curves. For clarity, only 13 of 40 neurons are shown. **B**,**E**: Density. **C**,**F**: Gain.

**Extended Data Fig. 3.**
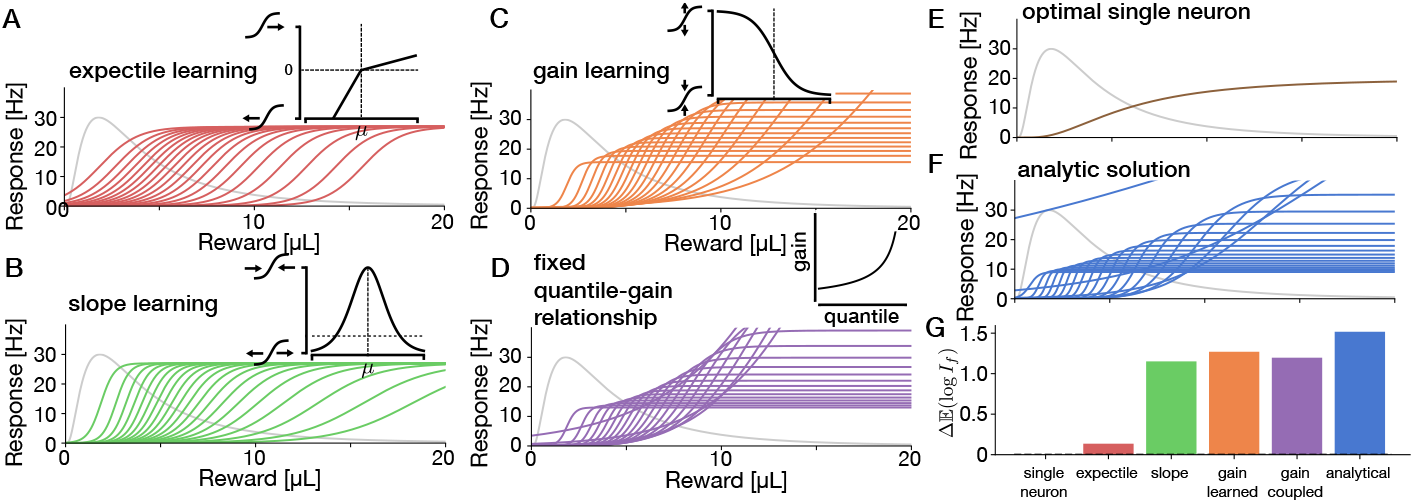
Evaluation of learning rules placing neurons’ midpoints at expectiles instead of quantiles. Plotting conventions as in Fig. 4. Each panel shows the converged population of 20 neurons after learning based on 20, 000 reward presentations. The inset illustrates the learning rule. **A**: Learning the position on the reward axis for the neurons to converge to the quantiles of the distribution. This learning rule is the distribution RL learning rule. **B**: Additionally learning the slope of the neurons to be proportional to the local density by increasing the slope when the reward falls within the dynamic range and decreasing otherwise. **C**: First method to set the gain: iterative adjustment to converge to a fixed average firing rate. **D**: Second method to set the gain: use a fixed gain per neuron based on the quantile it will eventually converge to. **E**: The efficient tuning curve for a single neuron. **F**: The analytically derived optimal solution. **G**: Comparison of information transfer across the different populations with the same number of neurons and expected firing rate.

## Supplementary Information

### Comparing thresholds and midpoints

**Fig. S1.**
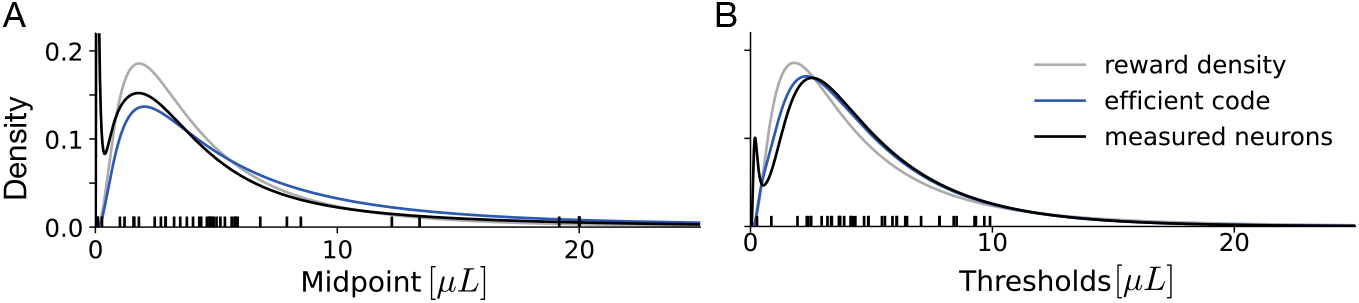
Log-normal kernel density estimation of midpoints and threshold. (**A**) and thresholds (**B**). Measured neurons (black) and efficient code (cyan) are overlayed over the reward density (gray).

Since we are using two metrics for the placement of neurons, threshold and midpoint, we discuss here how they compare. In our formalism, “density” refers to the midpoint of a neuron with a sigmoid response function. Threshold is the reward when a neuron fires at spontaneous activity; this is the point at which the neuron codes zero prediction error in reinforcement learning. Their distributions are slightly different. The relationship between the two depends on the relationship between the gain of the neuron (which defines the midpoint) and its spontaneous firing rate (which defines the threshold). A neuron whose gain exceeds twice its spontaneous firing rate has a higher midpoint than threshold, as reward needs to be higher for the firing rate to reach half the gain than to reach the spontaneous rate. The threshold will then lie in the lower, convex part of the response function. Correspondingly, a neuron with gain less than twice its spontaneous rate has a higher threshold than midpoint, and its threshold will lie in the upper, concave part of the response function. In the efficient code, gain increases with threshold. As a result, the prediction is that low thresholds are higher than their corresponding midpoints and high thresholds are lower than their corresponding midpoints. In turn, this means that the threshold distribution should be less dispersed than the midpoint distribution.

We illustrate this in the variable-magnitude task (Fig. S1). Empirically, the threshold distribution is indeed narrower (and taller) than the midpoint distribution, as predicted by the efficient code. Additionally, the spontaneous firing rate we estimated for this task is lower than the average half gain. Correspondingly, the average threshold (5.00 μL) is smaller than the average midpoint (5.96 μL). These shifts are all relatively small though, and the relationships with other variables (Fig. 2B, D, F) hold for midpoints as they do for thresholds.

## Footnotes

1 Ganguli and Simoncelli [18] used the interval [0, *N*], where *N* is the number of neurons, which is entirely equivalent.

## Notes

### Competing Interest Statement

The authors have declared no competing interest.

### Summary of Updates

This is a general overhaul of the paper for the next review step. A new task has been added, additional predictions have been added. All figures and results have been revised. A suggestion how the efficient code may be learned has been added.

